# Workflow for Assessing Mass Spectrometer Performance in Shotgun Lipidomics

**DOI:** 10.64898/2026.11.16.766365

**Authors:** Silvia Radrezza, HongKee Moon, Robert Ahrends, Britta Brügger, Cristina Coman, Christer S. Ejsing, Xianlin Han, Marcus Hoering, Saida Ibntaleb, Christian Klose, Jochen Koddebusch, Gerhard Liebisch, Marcellina Przybylska, Richard Sprenger, Julio L. Sampaio, Kai Schuhmann, Dominik Schwudke, Christoph Thiele, Hu Wang, Mohamed H. Yaghmour, Lena Hersemann, Andrej Shevchenko

**Affiliations:** Max Planck Institute of Molecular Cell Biology and Genetics, Dresden, Germany; SciComp, Max Planck Institute of Molecular Cell Biology and Genetics, Dresden, Germany; Institute of Analytical Chemistry, University of Vienna, Vienna, Austria; Heidelberg University Biochemistry Center (BZH), Heidelberg, Germany; Department of Biochemistry and Molecular Biology, Villum Center for Bioanalytical Sciences, University of Southern Denmark, Odense, Denmark; Barshop Institute for Longevity and Aging Studies at UT Health San Antonio, TX, USA; Institute of Clinical Chemistry and Laboratory Medicine, University Hospital Regensburg (UKR), Regensburg, Germany; Institut Curie, PSL Research University, CurieCoreTech Metabolomics and Lipidomics Platform, Paris, France; Lipotype GmbH, Dresden, Germany; Bioanalytical Chemistry, Research Center Borstel Leibniz Lung Center, Borstel, Germany; German Center for Infection Research (DZIF), Thematic Translational Unit Tuberculosis, Partner Site: Hamburg-Lübeck-Borstel-Riems, Borstel, Germany; Airway Research Center North, Member of the German Center for Lung Research (DZL), Site: Research Center Borstel, Borstel, Germany; Life & Medical Sciences Institute, University of Bonn, Bonn, Germany

**Keywords:** shotgun lipidomics, lipid quantification, Orbitrap mass spectrometers, high-resolution mass spectrometer, quality control, performance indices, lipid

## Abstract

Reproducible and testable analytical performance of mass spectrometers is critical for inter-laboratory concordance of quantitative lipidomics analyses. However, common instrument readout or analyses of arbitrarily chosen lipid mixtures do not lend themselves to direct comparison between instruments independently configured and operated in different laboratories under in-house settings. We developed and validated the QC Shotgun workflow and web-accessible software that enable rapid testing, evaluation, documentation, and longitudinal monitoring of the performance of high-resolution mass spectrometers in quantitative shotgun lipidomics. Completed in less than 5 minutes, the QC Shotgun test reports 8 relative metrics indices that facilitate inter-laboratory comparison and longitudinal assessment of instrument performance and identify deficiencies that are not apparent from instrument readouts. Systematic documentation of instrument status can improve inter-laboratory consistency and interpretation confidence of lipidomics datasets.

**Graphical abstract:** 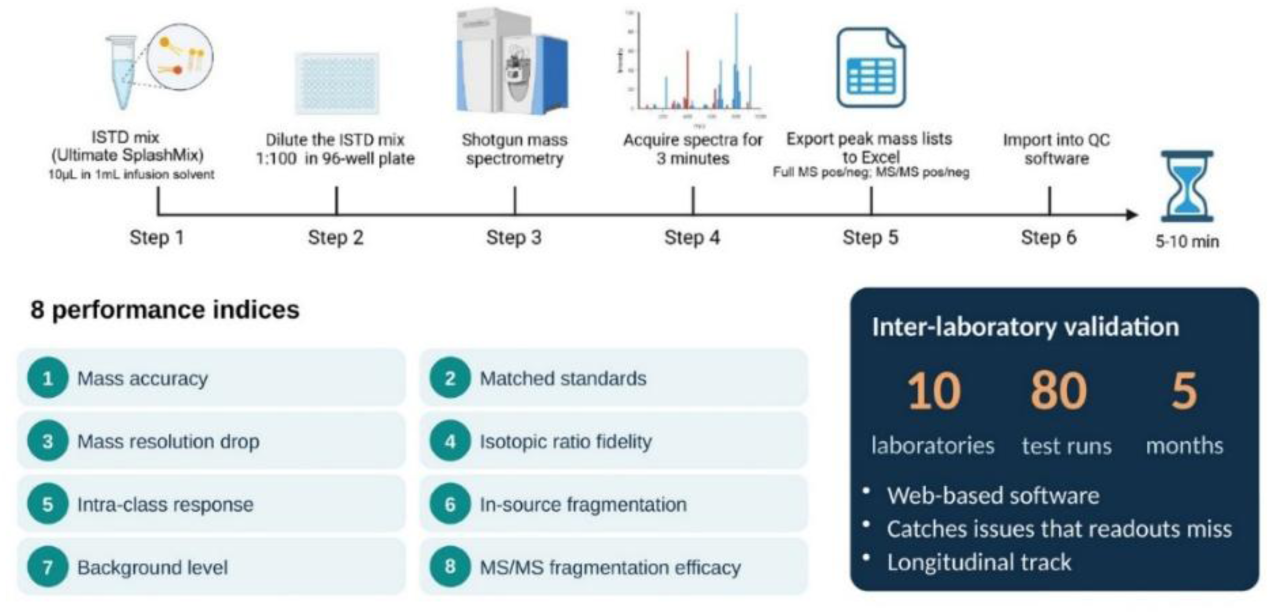

---

Within the past two decades, lipidomics has evolved from an exotic analytical tool used by a few lipid biochemistry aficionados to enabling technology supporting a broad spectrum of biomedical applications (reviewed in ^1^). This was timely recognised by lipidomics practitioners and prompted several community-driven efforts to develop field standards and operational protocols for the generation, processing, interpretation, reporting, and dissemination of lipidomics data ^2–6^.

The ring trials relied upon lipidome-wide quantification of the NIST 1950 human blood plasma standard ^7^ or targeted quantification of species of particular lipid classes (*e.g.* ceramides ^8^) and were instrumental for comparative evaluation of analytical methods across lipidomics laboratories. However, inter-laboratory discordance between lipid concentrations reported by study participants could be due to diverse technical factors, *e.g.* different lipid extraction protocols, employed standards, glass- and plasticware, operational settings of mass spectrometers and data processing software ^7^. Although lipid standards are widely available today, lipidome-wide quantification often employs one internal standard per lipid class and relies on assumptions that may or may not be fulfilled under particular experimental conditions ^9^.

It has come to our attention that the performance of mass spectrometers used for lipidomic analyses has received lesser attention. It is often presumed that, once a mass spectrometer is operational (*e.g.* all instruments readbacks are green) and there is no indication of a massive loss of the analyte signal, then the instrument is supposedly suitable for quantitative lipidomics. However, this assumption is not necessarily valid. Currently, there is no commonly accepted procedure for systematically evaluating and documenting mass spectrometers performance, despite its importance for longitudinal studies or high-throughput screens has been recognized.

Nowadays quantitative shotgun lipidomics most commonly relies on direct infusion of total lipid extracts into a high-resolution mass spectrometer (HR MS) with prior chromatographic separation of analytes (reviewed in ^10,11^). Lipids are identified by accurately measured masses ^12,13^ and/or characteristic fragment ions in MS^n^ spectra acquired from detectable precursors ^14^. The molecular specificity of shotgun identification benefits from high mass resolution and mass accuracy, dual spectra acquisition polarity ^15^, as well as long and stable spraying of the analyte at the micro- to nano-flow rate ^16^. Over years of routine lipidomics applications, shotgun lipidomics has demonstrated high inter-laboratory consistency of determined molar concentrations ^7,17^. However, factors contributing to its analytical robustness have not been fully understood.

We argue that the stable analytical performance of high-resolution mass spectrometers can be a major factor affecting lipidomics analyses and that it should be systematically checked, documented and, arguably, also reported. To this end, we set out to develop a technically simple testing workflow and assisting software to report composite performance indices informative for both longitudinal monitoring and inter-laboratory comparison. The workflow, supported by web-accessible QC Shotgun software, was used to evaluate and monitor the performance of Orbitrap mass spectrometers independently configured for shotgun lipidomics in 10 laboratories, each operating the instruments under their in-house settings. A longitudinal inter-laboratory study indicated high stability of key performance indices and was also instrumental for rapid troubleshooting and adjustment of deviating settings and conditions.

## MATERIALS AND METHODS

### Test sample

The exact protocol for preparing the mixture of lipid standards for the shotgun QC analysis was at the discretion of participating laboratories, and only the final dilution rate of 1:10000 of UltimateSPLASH™ ONE (Avanti Polar Lipids, cat. 330820L-1EA) was required. As a reference, we provide the protocol used in MPI CBG.

Aliquots of 10 µL of UltimateSPLASH™ ONE were prepared in Eppendorf tubes and stored at −20°C until use. Before testing, an aliquot was diluted to 1 mL by MSmix spraying solvent consisting of isopropanol/methanol/chloroform 4:2:1(v/v/v) supplemented with 7.5 mM ammonium formate. A further 1:100 dilution (1µL spiked into 99 µL of MSmix) was performed in the plate (Twin.tec 96-well PCR plate, skirted, colourless, 150 µL; Eppendorf SE, cat. 951020401) before MS analyses, which were performed in two technical replicates (10 µL sample loaded, of which 1.5µL of dead volume).

### Acquisition of test spectra

Shotgun spectra were acquired using local user-defined settings on hybrid quadrupole Orbitrap mass spectrometers equipped with ion sources available in participating laboratories. Below is the description of the acquisition protocol, ion source and instrument control settings used in MPI CBG.

FT MS and HCD FT MS/MS spectra were acquired on a Q Exactive mass spectrometer (Thermo Fisher Scientific, Bremen) equipped with a robotic nanoflow ion source, TriVersa NanoMate (Advion BioSciences, Ithaca, NY), using nanoelectrospray chips with a spraying nozzle diameter of 4.1 μm. Ionisation voltage was ±0.96 kV; gas backpressure was 1.25 psi in both modes; the temperature of the ion transfer capillary was 200 °C; S-lens RF level was 50%. Spectra were acquired in profile mode for a total time of 3 min. FT MS spectra were acquired within the range of *m/z* 150 to1000 from start to 1.0 min in positive mode and *m/z* 150 to1000 from 1.7 to 2.5 min in negative mode at the target mass resolution R _m/z_ =200 of 140.000 under automated gain control (AGC) value of 3 × 10^6^ and maximal injection time of 3000 ms. Parallel reaction monitoring (PRM) (time range of 1.0 to 1.25 min) was performed in positive mode by targeting PC-d5 33:1 (*m/z* 751.6008) and in negative mode (2.5 – 2.75 min) from PE-d5 33:1 (*m/z* 707.5393) precursor ions, respectively; precursor isolation window was 5.0 Th wide; nCE of 10% and 30%; AGC of 5 × 10^4^ with the maximum injection time of 3000 ms.

### Peak lists export

Raw files were opened in Xcalibur QualBrowser (Thermo Fisher Scientific), and a *ca.* 0.5 min time range of stable spray (as judged by the TIC trace) was selected separately for each acquisition polarity and FT MS or FT MS/MS mode. Peak lists containing *m/z*, intensity and resolution (see Dataset 1S as an example) were copied into a Microsoft Excel (.xlsx) file (one spreadsheet per modality in fixed order: FT MS positive, FT MS negative, FT MS/MS positive, FT MS/MS negative) and uploaded to QC Shotgun software.

### QC Shotgun software

QC Shotgun is a web-based quality control software developed using JavaScript and the Observable Framework HQ (2024) https://github.com/observablehq/framework.

QC Shotgun is accessible at https://shotgunqc.mpi-cbg.de/ and provides interactive visualisation, longitudinal tracking and exporting of performance indices in .csv format. Indices are not stored on the server and are deleted once the session is finished.

## RESULTS and DISCUSSION

### QC Workflow and Software for Shotgun Lipidomics

We proposed that comparative evaluation of the performance of high-resolution mass spectrometers in shotgun lipidomics could rely on a few composite relative metrics indices calculated from *m/z*, intensities and mass resolution of peaks of lipid standards detected in FT MS and HCD FT MS/MS spectra recognized by Xcalibur software (Thermo Fisher Scientific) supplied by the instrument manufacturer (**Figure 1**). Users could choose flow infusion rates, ionisation voltages or other settings that they optimised for their own ion source and mass spectrometer. It was only required *i)* to use UltimateSPLASH™ ONE lipid standards mix; *ii)* which was diluted at the fixed (1:10^4^, v/v) ratio by any appropriate spraying solvent containing mM concentration of ammonium formate; and *iii)* to acquire broadband FT MS spectra within a pre-defined *m/z* range in both polarities. For MS/MS evaluation, two lipid standards were fragmented in positive and negative mode under high (30%) and low (10%) nCE offset such that the peaks of the intact precursor and characteristic fragment ions were detected in the same MS/MS spectrum. Spectra acquisition time is arbitrary, although one minute of recording under stable (as judged by the TIC trace) spraying typically delivers good ion statistics. The four acquired spectra (FT MS and FT MS/MS in positive and negative modes) were opened and averaged using Xcalibur software; a *ca.* 30 sec time range with stable TIC was selected to produce four peak lists with *m/z*, raw intensity and mass resolution assigned by Xcalibur for each reported peak. Peak lists were saved as worksheets in a Microsoft Excel (.xlsx) file whose name included the acquisition date and test (or laboratory) name in “YYMMDD-[TEXT]” format. Finally, the .xlsx file was uploaded into the web-accessible QC Shotgun software, which computed performance indices and provided table and graphical output assisting their further consideration.

**Figure 1.**
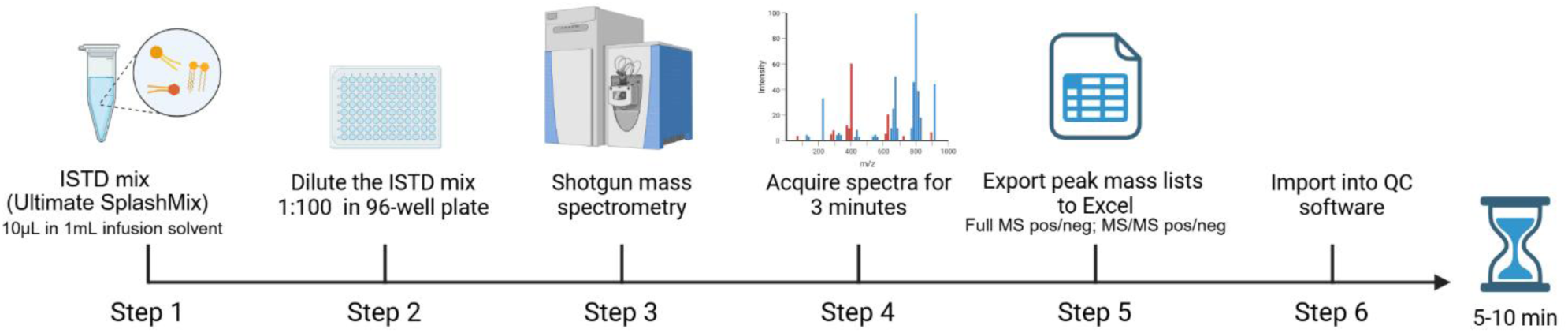
Workflow for testing the performance of a high-resolution mass spectrometer for shotgun lipidomics. It starts with direct infusion of an appropriately diluted mix of synthetic lipid standards with known molar concentrations into the mass spectrometer and acquisition of FT MS and FT MS/MS spectra in positive and negative modes, which usually takes *ca.* 3 min. Spectra are converted to peak lists using Xcalibur software and saved as worksheets in the same .xlsx file, which is then uploaded to the online QC Shotgun software. The software computes and reports, as text or graphical output, 8 composite performance indexes. Testing, including spectra acquisition and their evaluation by QC Shotgun, is usually completed in *ca.* 5 min.

In each peak list, QC Shotgun matched peaks *m/z* with expected masses of appropriate ion forms of lipid standards within ± 5 ppm mass tolerance. Next, *m/z*, raw intensities (taken as peak apex) and mass resolution of the matched peaks were used to calculate 8 composite indices reflecting mass spectrometer performance (**Figure 2**).

**Figure 2.**
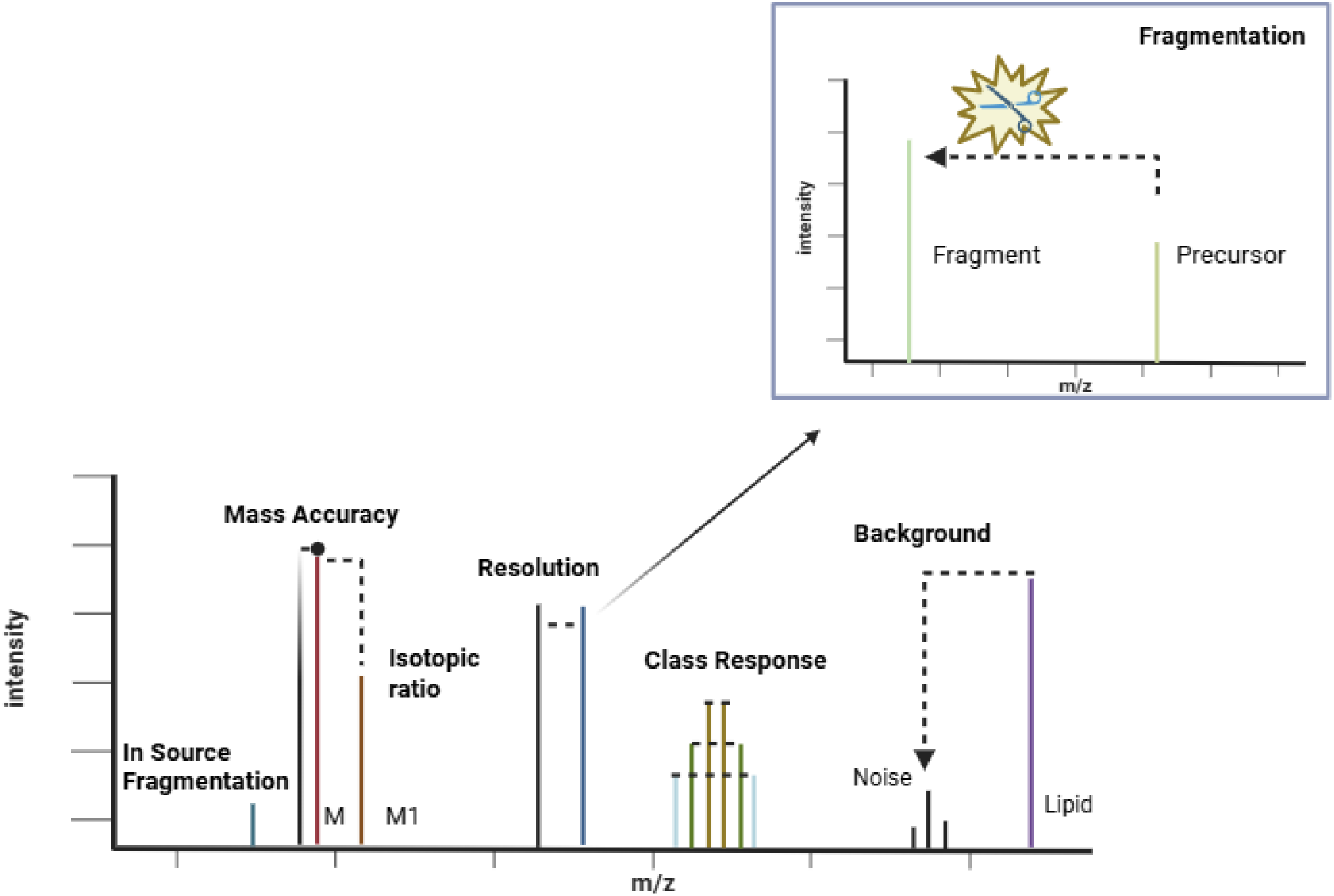
Schematic overview of performance indices computed by the QC Shotgun software. In each mode, the test acquisition produced one broad-band FT MS spectrum, from which QC Shotgun computed 7 relative indices. Mass accuracy, fidelity of isotopic ratios, mass resolution and its rate of decrease at higher *m/z* were computed directly from peak annotation. For other indices (in-source fragmentation; proportional response to the concentration of the same lipid class species; chemical noise level), abundances of different peaks were convoluted into a dimensionless relative value. Computing the index of MS/MS fragmentation efficacy used two separately acquired HCD FT MS/MS spectra. For details on index calculation, see text.

Below we explain in detail how the performance indices are computed.

#### Mass accuracy

It is the average value (in ppm) of mass differences between calculated and measured *m/z* of monoisotopic peaks of lipid standards matched within a tolerance range of 5 ppm in positive and in negative mode. The number of matched standards is reported in a separate column.

#### Mass resolution and rate of its m/z-dependent decrease

In Orbitrap mass spectrometers, mass resolution decreases with increasing *m/z* ^18,19^ and it is important to document the actual resolution at the start and end of a typical range of lipid masses. For convenience, QC Shotgun reports the resolution value for the low *m/z* standard and the percentage of resolution drop at the higher *m/z* relative to the lower *m/z* standard in each polarity (**Table 1**).

**Table 1.** Molecular ions of lipid standards used as reporters for calculating performance indices.

|  | **Positive** | **Negative** |
|---|---|---|
| Mass resolution drop | Cer-d7 36:2:2 (*m/z* 571.5790)<sup>^</sup><br>TG-d5 57:4 (*m/z* 947.8798)<sup>\$</sup> | LPC-d5 15:0 (*m/z* 531.3464)<sup>\*</sup><br>PC-d5 39:4 (*m/z* 873.6387)\* |
| Isotopic ratio concordance | TG-d5 49:1 (*m/z* 841.8007)<sup>\$</sup><br>Cer-d7 36:2:2 (*m/z* 571.5790)<sup>^</sup> | PC-d5 35:1 (*m/z* 823.6219)\*<br>LPC-d5 17:0 (*m/z* 559.3766)<sup>\*</sup> |
| Intra-class response | TG-d5 57:4 (*m/z* 947.8793)<sup>\$</sup><br>TG-d5 41:0 (*m/z* 731.6916)<sup>\$</sup><br>TG-d5 55:4 (*m/z* 919.8479)<sup>\$</sup><br>TG-d5 43:1 (*m/z* 757.7073)<sup>\$</sup><br>TG-d5 53:3 (*m/z* 893.8322)<sup>\$</sup><br>TG-d5 45:1 *(m/z* 785.7384)<sup>\$</sup><br>TG-d5 51:2 (*m/z* 867.8166)<sup>\$</sup><br>TG-d5 47:1 (*m/z* 813.7696)<sup>\$</sup><br>TG-d5 49:1 (*m/z* 841.8007)<sup>\$</sup> | PE-d5 31:1 (*m/z* 679.5097)<sup>#</sup><br>PE-d5 39:4 (*m/z* 785.5887)<sup>#</sup><br>PE-d5 33:1 (*m/z* 707.5415)<sup>#</sup><br>PE-d5 37:3 (*m/z* 759.5727)<sup>#</sup><br>PE-d5 35:1 (*m/z* 735.5730)<sup>#</sup> |
| In-source fragmentation | PC head group *(m/z* 184.0736) | FA 17:0 (*m/z* 269.2486) |
| MS/MS efficacy | PC head group (*m/z* 184.0736) | FA 16:1 (*m/z* 253.217) |

*Footnote:* Symbols in superscript indicate the molecular form of detected lipid precursors: ^ for [M+H]<sup>+</sup> <sup>\$</sup> for [M+NH<sub>4</sub>]<sup>+</sup>; <sup>#</sup> for [M-H]<sup>-</sup> and  \* for [M+HCOO]<sup>-</sup>

#### Concordance of isotopic ratios

Accurate representation of isotopic profiles is important for shotgun quantification ^20,21^, but it can be affected by several operational settings such as AGC or IT ^16^. To evaluate the fidelity of isotopic profiles, QC Shotgun calculates the ratio (in %) of theoretical to measured intensity of the first isotopic to monoisotopic peaks for two standards separately in positive and in negative mode (**Table 1**). For each polarity the average value of the two ratios is reported; the value of 100% indicates a perfect match between measured and expected intensity ratios.

#### Proportional peak intensity for the same class lipid species

In shotgun analysis, the analyte composition should not change over time and the intensity of peaks of the same lipid class standards is expected to be proportional to their molar concentration ^9,22^. As a measure of unbiased instrument response, QC Shotgun computes the relative standard deviation (RSD) of peak intensities of nine triacylglycerol and of five phosphatidylethanolamine standards in positive and negative mode, respectively. The intensity of their monoisotopic peaks is first adjusted to the number of carbon atoms in each lipid and then normalised to its concentration as provided by the manufacturer (**Table 1**). RSD value of zero indicates no difference between normalised peak intensities.

#### Impact of in-source fragmentation

In-source fragmentation can degrade molecular ions of lipids already during analyte ionisation and bias the lipidome quantification. QC Shotgun assesses the impact of in-source fragmentation by calculating the ratio between the intensity of a characteristic fragment ion and the total intensity of all of its potential precursors in the FT MS (not in FT MS/MS) spectrum. In-source fragmentation of [M+H]^+^ precursors of PC and SM yields the phosphorylcholine fragment with *m/z* 184.0736. QC Shotgun calculates its ratio to the sum of intensities of five PC, three LPC and five SM standards. In negative mode, QC Shotgun calculates the ratio of the intensity of the acyl anion of FA 17:0 (*m/z* 269.2486) to the sum of intensities of [M-H]^−^ions of PE, PG, PI and all lyso-lipids containing the FA 17:0 moiety. A zero value indicates that no in-source fragmentation occurs.

#### Relative intensity of background peaks

Abundant peaks of diverse chemical contaminants affect the specificity and sensitivity of shotgun analysis. QC Shotgun estimates the level of background irrespective of its molecular origin by reporting the ratio of the summed intensities of all peaks of lipid standards (including their 1^st^ and 2^nd^ isotopes) to all other peaks in the FT MS spectrum. A higher value indicates lower background and presumably higher overall sensitivity.

#### MS/MS fragmentation efficacy

Several instrument-dependent factors (*e.g*. insufficient collision gas pressure or excessive fragment ions scattering due to ion optics contamination) can affect the performance of FT MS/MS. To assess HCD fragmentation efficacy, QC Shotgun computes the ratio of intensity of precursor ions acquired at low collision energy (nCE 10%) and of the characteristic fragment acquired at higher (nCE 30%) collision energy using a stepped collision energy acquisition. Lower ratios indicate reduced sensitivity in HCD FT MS/MS.

Taken together, these 8 indices capture key features of high-resolution mass spectrometers essential for shotgun lipidomics. The indices either have a direct physical meaning (*e.g.* mass accuracy) or are relative measures computed from peak intensity ratios, rather than absolute intensities of the peaks that may vary between experiments. Therefore, they could be applicable for inter-laboratory comparisons.

### Benchmarking of QC Shotgun Workflow

We next checked whether these composite indexes adequately reflected the instrument status and could predict its performance in shotgun analyses. To this end, we deliberately changed instrument settings and spectra acquisition conditions to suboptimal values, performed the QC Shotgun test, and checked if and how the indices respond to compromised instrument performance.

We first checked how QC Shotgun test responded to sub-optimal AGC values (Dataset 2S). The normal AGC target value for the hybrid QExactive Orbitrap mass spectrometer in MPI CBG is 5 × 10^4^. Setting AGC to lower (2 × 10^4^) value did not affect mass resolution and accuracy, but worsened isotopic ratio fidelity because of limited ion statistics. Increasing AGC to maximal allowed value of 5 × 10^6^ also did not impact the resolution, but severely compromised mass accuracy, consistent with enhanced ions coalescence ^23^ (Figure 3S). QC Shotgun reported that masses of *ca* 20% of lipid standards in positive and of 85% in negative mode fell outside the permitted 5 ppm error range (Dataset 2S) such that no indices relying on multiple standards (Table 1) could be computed. High AGC impaired both isotopic ratios fidelity and the proportionality between peak intensities and standard concentrations.

**Figure 3.**
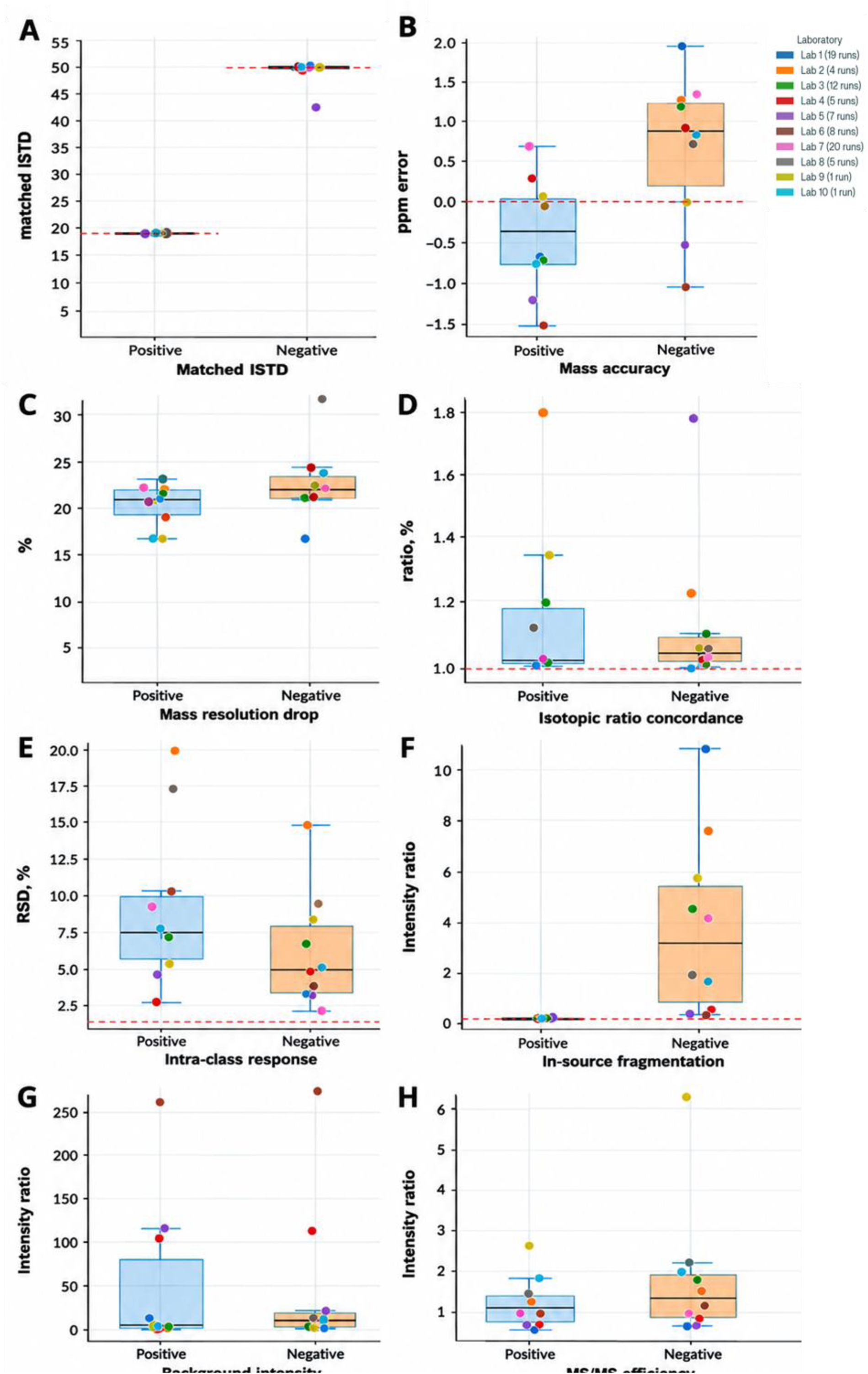
Inter-laboratory comparison of median values of performance indices. Indices were computed from all QC measurements submitted by each participating laboratory. Inset: color coding of laboratories and the number of submitted QC tests. Red dotted lines: the best expected value for a particular index (if available); *e.g.* the lowest possible mass error is zero. Units in other panels are explained in the text. Panels with performance indices: A, number of matched lipid standards; B, mass error; C, drop in mass resolution, %; D, fidelity of isotopic ratios, arbitrary units, a.u.; E, proportional peak intensity, %; F, in-source fragmentation, a.u.; G, ratio of lipid peaks to background, a.u.; MS/MS fragmentation efficiency, a.u.

Disabling the lock mass function in freshly calibrated mass spectrometer did not measurably affect its performance. Interestingly, increasing the ion transfer capillary temperature from typical 200 °C to 300 °C did not enhance in-source fragmentation, but improved the proportionality between peak intensity and concentration of lipid standards and reduced background (Dataset 2S). We note, however, that this might not hold true for unstable lipids.

The QC workflow also helped to pinpoint inadvertent errors in analyte preparation. When ammonium acetate was accidentally used instead of required ammonium formate, QC Shotgun reported fewer matched standards, while other indexes were unaffected (Dataset 2S). Occasional contamination of the spraying solvent with a PEG-like polymer likely originating from an untested brand of plasticware (Figure 1S) did not affect resolution, mass accuracy or the number of matched standards. However, it enhanced spectrum background and worsened both isotopic ratios and proportionality of peak intensities (Dataset 2S).

We therefore concluded that QC Shotgun test adequately reflected sub-optimal instrument performance resulting from incorrect settings or analyte contamination. These experiments also reinforced the notion that the diagnostics should not rely on a single easy measurable parameter such as mass deviation of a reference peak or mass resolution reported by some basis software. Instead, its assessment should incorporate a few independently computed indices reflecting different aspects of the instrument analytical performance.

### Inter-laboratory longitudinal comparison of performance indexes

To conduct the longitudinal trial, 10 laboratories applied the QC workflow to their Orbitrap family mass spectrometers configured for shotgun lipidomics. An aliquot of the UltimateSPLASH standard was diluted on site, and spectra were acquired using in-house laboratory settings (*e.g.* flow rate of analyte infusion, ionisation voltage and the composition of spraying solvent) several times within the period of 5 months. The laboratories reported if and when the calibration or repair of their instruments was carried out. Peak lists were produced locally and processed by QC Shotgun software in MPI CBG. Altogether, 80 test runs were collected (Dataset 2S), with the longest sequence of 20 runs collected by the same laboratory. The input from individual laboratories was anonymised while the original submission dates were retained; raw data were not shared and were not subjected to back-checking and correction. **Figure 3** exemplifies the concordance between median values of the 8 performance indices calculated from the input of individual laboratories over the entire test period.

Altogether, comparative analysis demonstrated good concordance of performance indices between laboratories. Mass accuracy was routinely better than ± 2 ppm throughout the testing period, while median values calculated for the full set of lipid standards were below ± 1 ppm in both modes (**Figure 3A**, **3B**).

The Orbitrap firmware sets the target mass resolution. However, the decline in mass resolution at higher *m/z* varied between instruments by 5 to 10 % in negative and positive modes, respectively (excluding two notable outliers) (**Figure 3C**). With the exception of a few outlying measurements, the Orbitrap instruments accurately determined isotopic ratios, although the intensity of the first isotopic peak was consistently lower than expected by *ca.* 10% in negative and 20% in positive mode (**Figure 3D**).

Proportionality between peak intensities and concentrations of the same class lipids is a key assumption underlying absolute quantification by FT MS while it typically relies on a few or just one standard per each class. Comparing median values suggested that RSD between normalised intensity of lipid peaks was below 20%, and for five laboratories that submitted the most complete datasets it was < 10% in both polarities (**Figure 3E**). We argue that, effectively, this sets the level of accuracy for absolute (molar) quantification achievable by shotgun analyses on Orbitrap instruments.

Under normal operational settings, the impact of in-source fragmentation for common lipid classes (*e.g.* glycero- and glycerophospholipids; ceramides) was almost negligible in positive mode, but was more pronounced in negative mode, as judged by a 3.5-fold (median) higher ratio of intensity of the reporting fragment to summed intensities of its possible precursors (**Figure 3F**). However, this difference may also reflect our choice of carboxylate anion of 17:0 fatty acid as the reporting fragment ion, which might occasionally originate from solvent or plastic contaminant and, if in doubt, should be checked in blank run.

We operationally defined the level of chemical noise as a ratio of intensities of all lipid peaks to all other peaks in the spectrum. While it was stable across labs and individual measurements, three laboratories (laboratories 4, 5, 6) achieved superior signal-to-noise ratio in both positive and negative modes (**Figure 3G**). However, laboratories 5 and 6 chose to infuse the analyte at 10x and 100x higher concentrations than other laboratories and corresponding index values could not be directly compared.

MS/MS fragmentation efficacy test is meant to discover accidental problems with the instrument set-up that might affect fragment ion intensity, *e.g.* insufficient collision gas flow or malfunctioning electronics, rather than assisting in fine-tuning of instruments. The index values were generally consistent across the full dataset (**Figure 3H**).

Effectively, median values of indices defined a consensus inter-laboratory reference of instruments performance. By comparing actual and consensus index values users could spot “acute” problems such as contaminated solvent or sensitivity loss. It is also helpful for detecting negative trends that require re-calibration and thorough inspection of instruments–which may not be apparent from a quick check on mass accuracy or resolution. Together with acceptable mass accuracy, tests in laboratory 2 revealed below the average isotopic concordance, intensity vs concentration proportionality and elevated level of in-source fragmentation (**Figure 3 D-F**), which indicate that current settings are seemingly sub-optimal for “generic” shotgun analyses.

However, median values do not characterise the robustness of instrument performance over an extended period of time, especially if it included its maintenance and calibration. To this end, we selected indices that are affected by both instrument and ion source performance and compared the magnitude of their variation between the laboratories (**Figure 4**).

**Figure 4.**
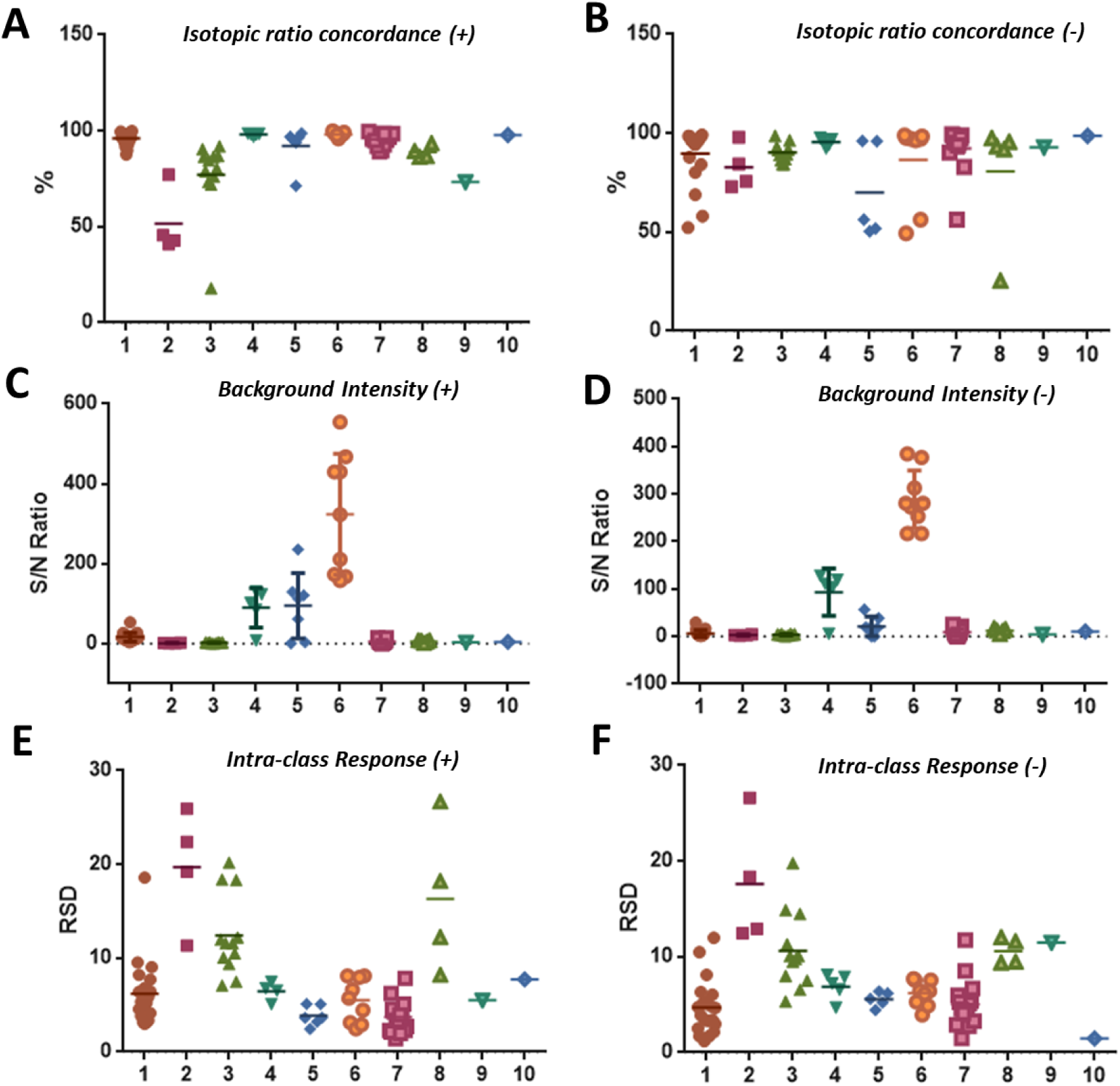
Variation of performance indices values. Panels A and B – fidelity of isotopic ratios; C and D: ratio of lipid peaks to background; E and F: proportional peak intensity. X-axes: the identification number assigned to each participating laboratory (as in Figure 3 and Dataset 3S); Y-axes: index value; units are the same as in Figure 3. Each datapoint represents individual measurement. Short horizontal lines indicate the median value of each index for each laboratory.

We observed that, particularly for the laboratories that submitted data from more than 10 time points, individual measurements clustered around the median, with the spread of < 15% and also result in similar inter-laboratory median values (**Figures 3 and 4**). Values from less frequent measurements may vary significantly (by 50% or even higher), particularly for isotopic ratios fidelity and proportional response indices, which may lead to biased lipidomics measurements. While a single measurement may help to identify acute problems with instrumentation set-up by comparing indices to the reference values, it also highlights the importance of consistent longitudinal measurements.

QC Shotgun software supports accumulation and graphical presentation of longitudinal trends of all indices. To illustrate the use of longitudinal profiles, we selected one of the largest available datasets of 19 test runs acquired by laboratory 1 within ca. 6 months. We further plotted the mass error (as the most stable) and proportional peak intensity (as most variable) indices (**Figure 5**). In this series of measurements, mass error was systematically offset (median ± SD) to lower masses – 0.7 ppm ±0.5 ppm in positive and to higher masses +2.0 ppm ± 0.7 ppm in negative mode with respect to the calculated lipid masses. Except single time point the mass difference did not exceed 3 ppm.

**Figure 5.**
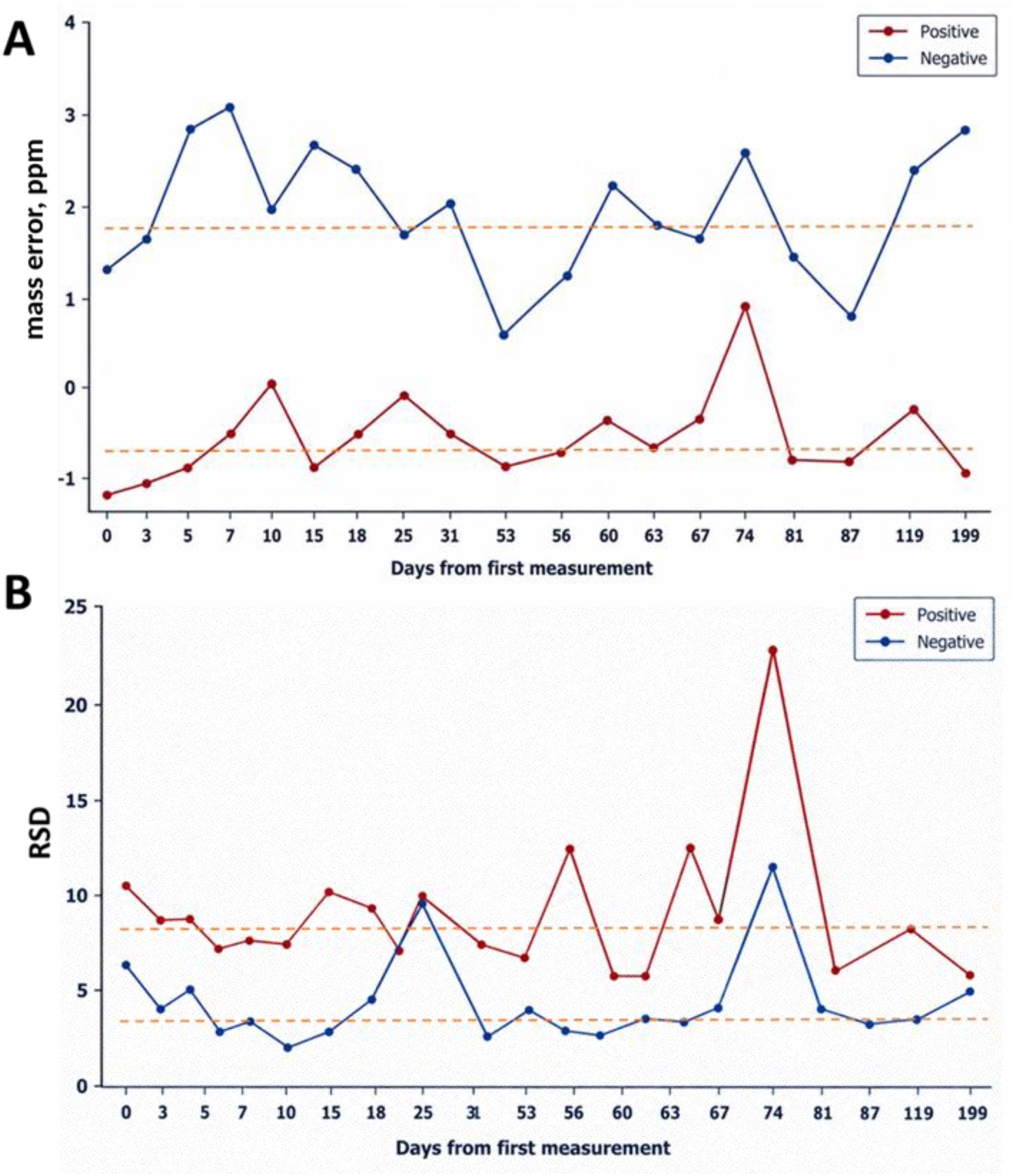
Longitudinal monitoring of mass error (panel A) and proportional peak intensity (panel B) acquired by laboratory 1. The best possible value for both indices is zero. For presentation clarity, the time scale on the x-axis is not linear; median values are indicated by dashed lines.

Within the range of 60 days, the intensities proportionality index was stable (mean ± SD) both in positive (8.7% ±4.2%) and in negative (4.2% ±2.6%). However, after 74 days the value of proportionality index more than doubled and this was also accompanied by elevated mass error clearly visible in the positive-mode trendline.

Taken together, longitudinal reports generated by QC Shotgun could be used in at least three ways. First, median values and their variance could benchmark the instrument performance against other laboratories employing shotgun lipidomics technology. Second, sudden changes in values of several indices could indicate (but not diagnose) acute instrumentation problems and prevent analyzing samples under sub-optimal conditions. This is particularly important for large-scale screens, in which data are typically analysed once the entire dataset has been acquired. Third, tracking index dynamics over time can reveal persistently accumulating performance deficiencies, while helping to avoid false alarms triggered by insignificant or occasional deviations in index values.

## CONCLUSIONS AND OUTLOOK

QC Shotgun workflow and software enable rapid systematic testing of the performance of high-resolution (here – Orbitrap) instruments in quantitative shotgun lipidomics applications. Eight performance indices reported in relative metrics and support longitudinal inter-laboratory comparison. The indices reflect different analytical features (*e.g.* mass accuracy, resolution, proportionality between peak intensity and concentrations, *etc.*) critical for shotgun lipidomics that could be affected by multiple instrumentation- and analyte-dependent factors. Median values of performance indices were relatively stable across ten participating laboratories and could be used as a reference for evaluating actual performance of high-resolution mass spectrometer.

By repetitive longitudinal measurements, QC Shotgun could create an objective performance history that helped to distinguish temporary deviations from emerging technical problems. We reasoned that the output of systematic testing could be converted into some kind of “black-box” record and included into raw datasets. Obviously, this does not warrant accurate lipidomic analyses, yet it provides a useful indication that the mass spectrometer was correctly configured and tuned. It might also help to alert operators to excessive chemical noise and associated reduction in sensitivity, increased rate of false positive assignments, compromised quantitation capacity, in-source fragmentation of “fragile” lipids or lack of MS/MS sensitivity. Fluctuations in performance indices can be promptly spotted and prompt the appropriate interventions (*e.g.* calibration, service, replacing chemicals or plasticware, *etc.*).

Here we demonstrated that the performance tests could be rapid, technically straightforward and produce a few composite indices reported in the format of relative units. QC tests are not meant to be truly diagnostic; rather, they should alert operators to emerging or persisting instrumentation problems. In contrast analysing complex samples (*e.g.* a standard sample of human plasma) is considerably more laborious and prone to uncertain interpretation, and is more appropriate for comprehensive ring trials across many participating laboratories.

QC Shotgun workflow only supports direct infusion lipidomics. However, given community interest, it should be possible to develop a convenient protocol also for LC-MS/MS based lipidomics, in line with QC routines already available in proteomics ^24^.

## ABBREVIATIONS

AGC: Automatic Gain Control
FA: Fatty Acid
FT MS: Fourier-Transform Mass Spectrometry
HCD: Higher Energy Collision Dissociation
HRMS: High Resolution Mass Spectrometry
IT: Injection time
LPC: Lysophosphatidylcholine
nCE: normalised Collision Energy
PC: Phosphatidylcholine
PE: Phosphatidylethanolamine
PG: Phosphatidylglycerol
PI: Phosphatidylinositol
QC: Quality Control
RSD: Relative Standard Deviation
SM: Sphingomyelin
TIC: Total Ion Current

## ASSOCIATED CONTENT

### Supporting Information

The Supporting Information is available free of charge at:

Figures showing TIC of test runs and extended data quality assessment (PDF). Dataset with the example of peak lists for input into QC Shotgun software; extended dataset with QC Shotgun results of test runs under manipulated instrument settings; QC Shotgun results of all tests in all participating laboratories (XLSX).

## AUTHOR INFORMATION

### Author contributions

SR conceived the study, validated the software, processed results and wrote the manuscript; HKM developed software; RA, BB, CC, CSE, XH, MH, SI, CK, JK, GL, MP, RS, JLS, KS, DS, CT, HW, MHY carried out test runs in participating laboratories; LH supervised software development; AS conceived and conceptualised the study, acquired funding and wrote the manuscript.

### Notes

The authors declare no competing financial interest.

## ACKNOWLEDGMENTS

We are grateful for members of the AS group in MPI CBG for expert support and discussion; Computer Department and Scientific Computing Facility of MPI CBG for expert support for web hosting of QC Shotgun software.

This work has been supported by the following funders: LiSyM DEEP-HCC from BMBF and CRC412 from DFG for AS; German Network for Bioinformatics Infrastructure (de.NBI, W-de.NBI-006) for DS; VILLUM Center for Bioanalytical Sciences (VKR023179) for CSE; is funded by the Deutsche Forschungsgemeinschaft (DFG, German Research Foundation) - Project Numbers 511488495 SFB - 1638/1 (Z02) and 278001972 - TRR 186 (Z04) data storage service SDS@hd supported by the Ministry of Science, Research and the Arts Baden-Württemberg (MWK) and the German Research Foundation (DFG) through grant INST 35/1803-1 FUGG and INST 35/1804-1 LAGG for BB.

