## Supporting Information for "Workflow for Assessing Mass Spectrometer Performance in Shotgun Lipidomics"

**List of Content:**

**Figure 1S.** TIC and FT MS spectra if clean and PEG-contaminated samples.

**Figure 2S.** Assessment of MS data quality across particiapting labs.

**Figure 3S.** AGC-dependent mass shift of internal standards in negative mode FT MS spectra.

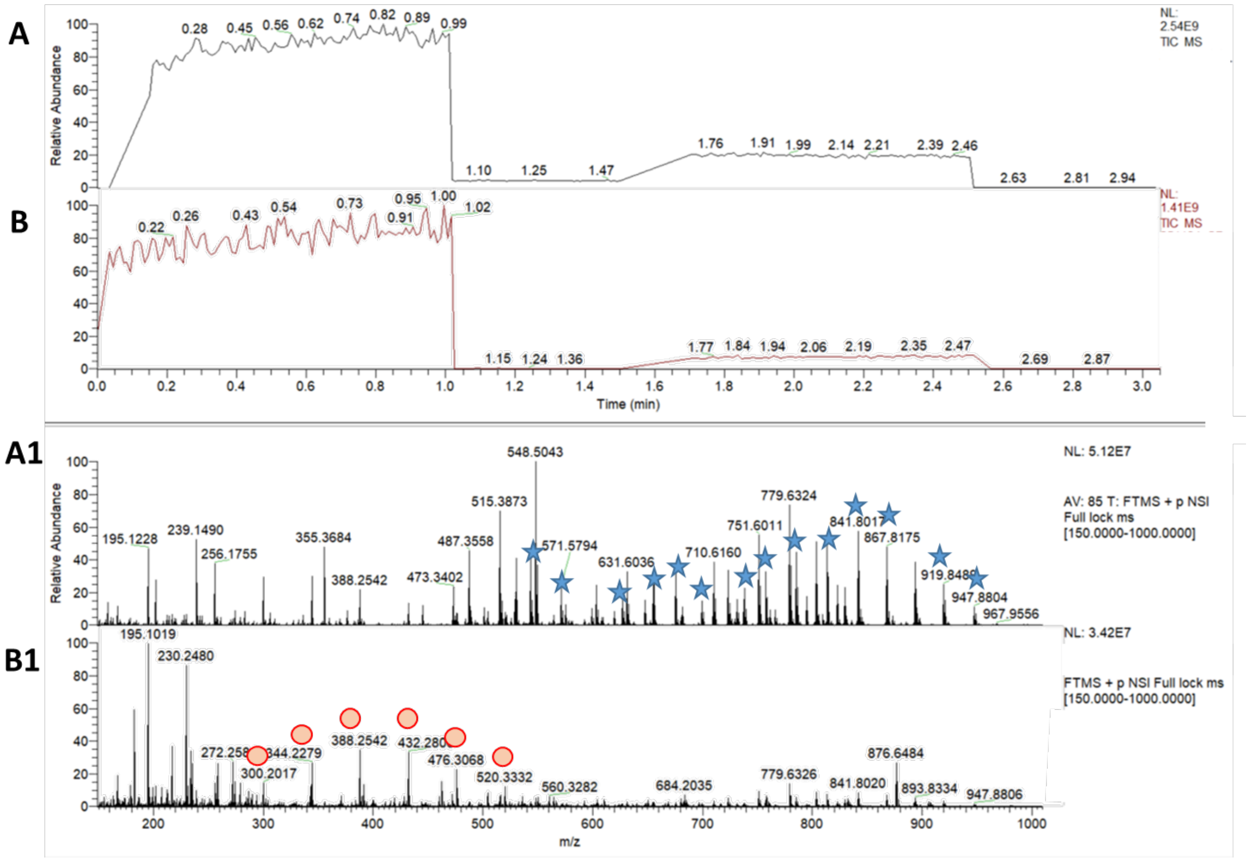


**Figure 1S. TIC and FT MS spectra of clean and PEG-contaminated samples.**

Total ion chromatograms (TIC) and full-scan MS spectra comparing clean versus PEG-contaminated spectra, demonstrating ISTD clear detection versus polymer interference. (A) TIC and full MS of clean extract. (A1) Extracted ion chromatograms show clear detection of internal standards (flagged) without zoom, confirming method sensitivity. (B) TIC and full MS of PEG-contaminated extract. (B1) Dominant PEG oligomer cluster ($m/z$ = 44 Da; $[PEG+n(C_2H_4O)+cation]^+$ series) suppresses ISTD signals (not visible even at full scale), illustrating contamination impact on quantitative lipidomics.

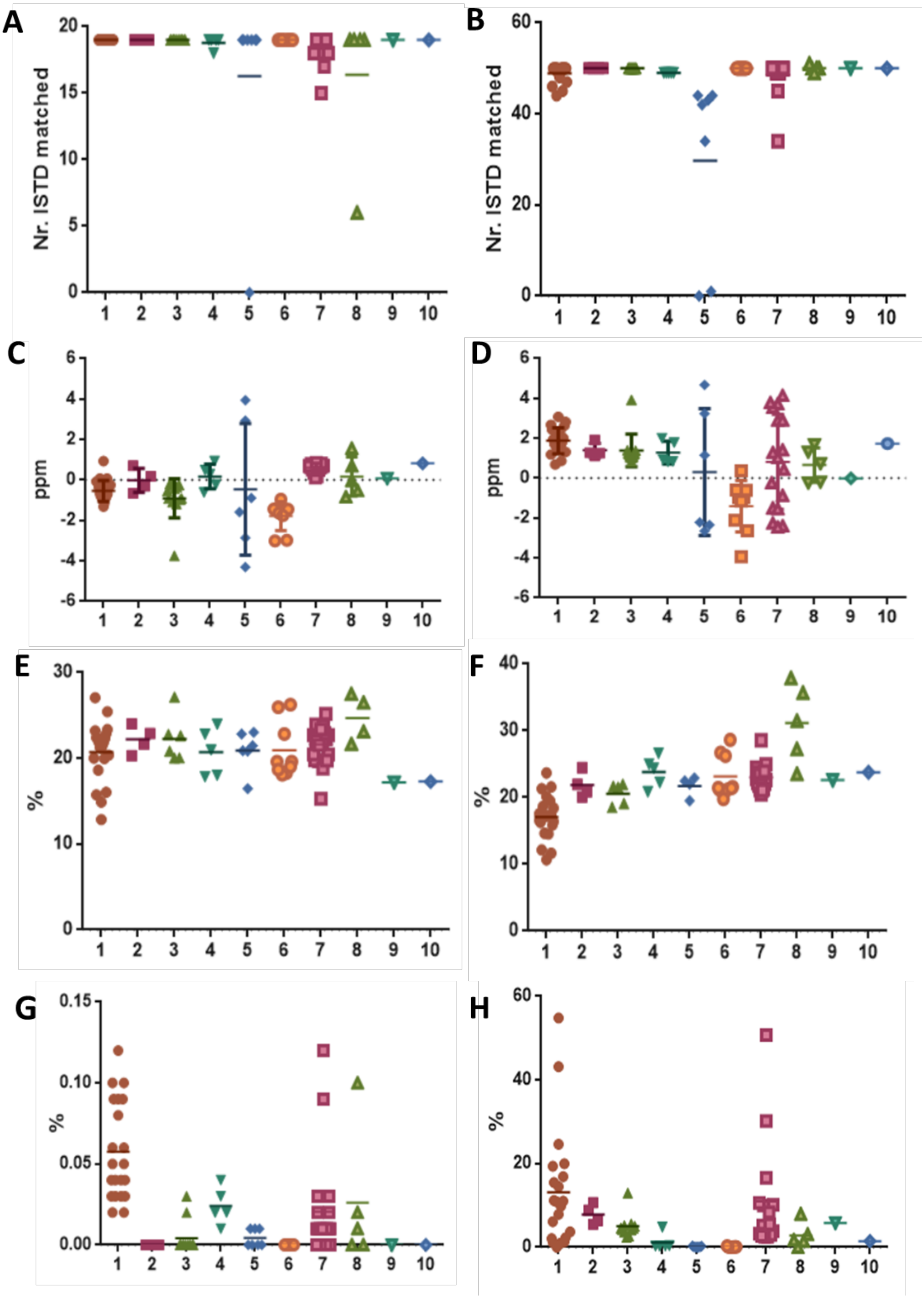
A
Nr. ISTD matched
0
5
10
15
20
1 2 3 4 5 6 7 8 9 10
B
Nr. ISTD matched
0
20
40
60
1 2 3 4 5 6 7 8 9 10
C
ppm
-6
-4
-2
0
2
4
6
1 2 3 4 5 6 7 8 9 10
D
ppm
-6
-4
-2
0
2
4
6
1 2 3 4 5 6 7 8 9 10
E
%
0
10
20
30
1 2 3 4 5 6 7 8 9 10
F
%
0
10
20
30
40
1 2 3 4 5 6 7 8 9 10
G
%
0.00
0.05
0.10
0.15
1 2 3 4 5 6 7 8 9 10
H
%
0
20
40
60
1 2 3 4 5 6 7 8 9 10

**Figure 2S. Assessment of MS data quality across participating laboratories.** A) Number of ISTDs identified in positive mode and B) negative mode by each laboratory (on the x-axis) across the measurements; C) distribution of mass accuracy error (in ppm) expressed as average for each acquisition in positive and D) negative mode; E) Mass resolution drop (%) for each measurement in positive and F) negative; G) In-source fragmentation in positive and H) negative, expressed in %; I) MS/MS fragmentation between precursor and fragment in positive and L) negative

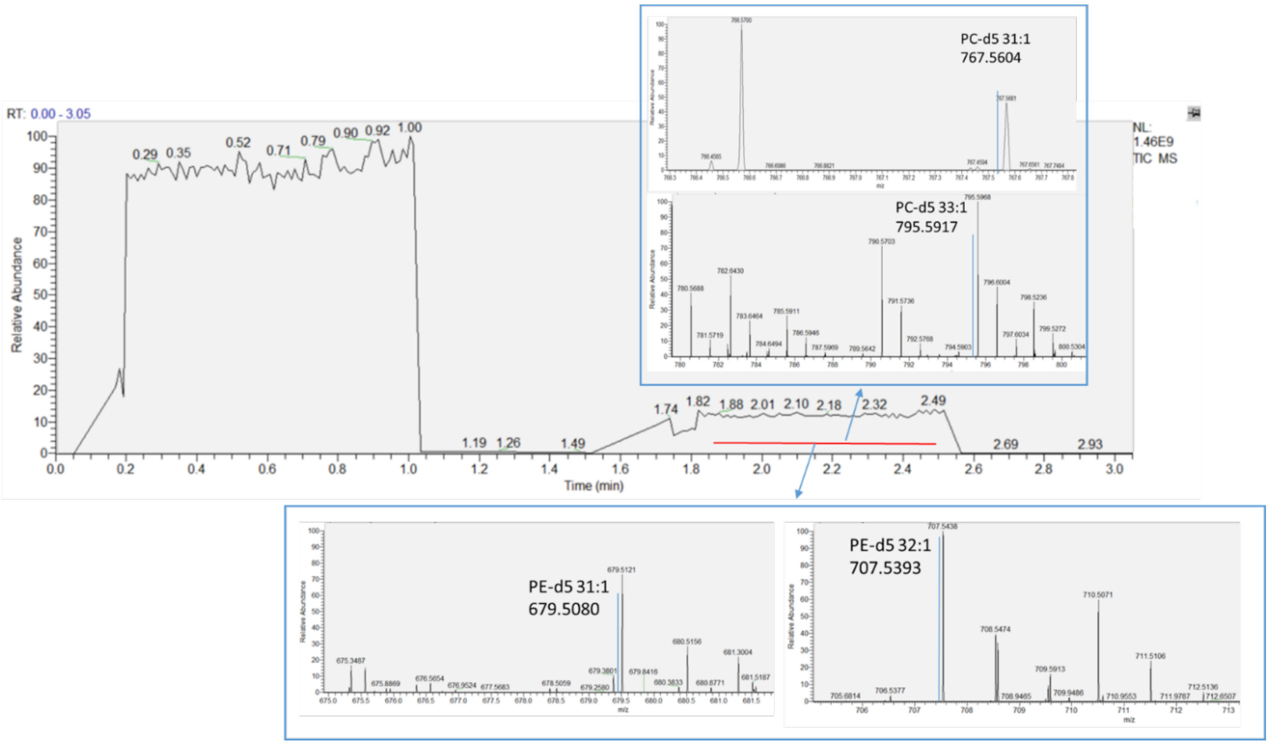


**Figure 3S. AGC-dependent mass shift of internal standards in negative mode FT MS spectra.**

Mass shift induced by high AGC on ISTDs in negative Full MS mode.
